# CatSper mediates the chemotactic behavior and motility of the ascidian sperm

**DOI:** 10.1101/2021.07.06.451356

**Authors:** Taiga Kijima, Daisuke Kurokawa, Yasunori Sasakura, Michio Ogasawara, Satoe Aratake, Kaoru Yoshida, Manabu Yoshida

## Abstract

Sperm motility, including chemotactic behavior, is regulated by changes in the intracellular Ca^2+^ concentration. The cation channel of sperm (CatSper), plays an important role in the regulation of intracellular Ca^2+^ concentration. In mammals, CatSper is the only Ca^2+^ channel that functions in the sperm, and the mice that lack the genes for the subunits of CatSper, which make up the pore region of the Ca^2+^ channel, are infertile due to the inhibition of hyperactivation of the sperm. CatSper is also thought to be involved in chemotaxis in sea urchins. In contrast, in the ascidian, *Ciona intestinalis*, the sperm-activating and -attracting factor (SAAF) interacts with Ca^2+^/ATPase, which is a Ca^2+^-pump. Although the existence of *CatSper* genes has been reported, it is not clear whether CatSper is the specific Ca^2+^ channel that functions in the ascidian sperm. Therefore, in this study, we generated *Catsper3* knockout (KO) animals that found that they were significantly less motile, with few motile sperms not exhibiting any chemotactic behavior. These results suggest that CatSper plays important roles in the spermatogenesis and basic motility mechanisms of sperms in both ascidians and mammals.

## Introduction

Ca^2+^ plays important roles in numerous cellular events. In the process of fertilization, Ca^2+^ plays indispensable roles by controlling many sperm functions, including the activation of sperm motility, sperm chemotaxis, capacitation and hyperactivation of motility, and the acrosome reaction (1,2). In particular, Ca^2+^ regulates the flagellar beating patterns (3,4) and the sperm swimming direction (5). The concentration of intracellular Ca^2+^ ([Ca^2+^]_i_) is regulated by Ca^2+^ influx via Ca^2+^ channels and Ca^2+^ efflux driven by Ca^2+^ exchangers or Ca^2+^ pumps; therefore, both Ca^2+^ channels and Ca^2+^ pumps play important roles in the sperm functions.

In ascidians, an increase in [Ca^2+^]_i_ also influences the sperm motility and chemotactic behavior towards the egg (5–7). The sulfate-conjugated hydroxysteroid sperm-activating and -attracting factor (SAAF) acts as a sperm activator and attractant in ascidians (8,9). The spermatozoa showing chemotactic behavior exhibit transient increase in [Ca^2+^]_i_ in the flagella (Ca^2+^ bursts), when the concentration of SAAF decreases to a local minimum, forming an asymmetric flagellar waveform of the sperm and inducing a series of sperm movements, consisting of turning and straight swimming (5,6,10). Regulation of [Ca^2+^]_i_ by SAAF is mainly achieved by controlling the efflux of Ca^2+^, which binds to and activates Ca^2+^/ATPase (PMCA) on the sperm plasma membrane, which is a Ca^2+^ pump responsible for the Ca^2+^ efflux (11). However, the specific molecules involved in the influx of Ca^2+^ remain unknown, although the involvement of store-operated Ca^2+^ channels has been suggested in pharmacological studies (6).

Among many Ca^2+^-regulating systems, the Ca^2+^ channel on the plasma membrane is an important player for the influx of Ca^2+^. In the mammalian sperm, the cation channel of sperm (CatSper), which is the sperm-specific Ca^2+^ channel, plays a crucial role in the regulation of sperm function. CatSper is a channel that shares homology with a voltage-gated Ca^2+^ channel (12), consisting of four pore-forming subunits (CatSper 1, 2, 3, 4) (13–15) and several auxiliary subunits (CatSper β, γ, δ, ε, ζ, and EFCAB9) (16–19). CatSper is located in the sperm flagella and mediates hyperactivation (12,15,20,21). Furthermore, CatSper seems to be the only Ca^2+^ channel that works in mammalian sperm (22) therefore, it is considered to mediate not only hyperactivation, but also the flagellar beating and swimming patterns (23).

In contrast to its crucial role in mammals, CatSper is absent in many animals including vertebrates: birds, amphibians, and many teleostean fishes that lack the *CatSper* gene (24). Furthermore, the expression levels and functions of CatSper proteins in animals, other than mammals, have been poorly investigated; however, CatSper appears to regulate the sperm chemotaxis in sea urchins (25). Moreover, the genome database of the ascidian, *Ciona intestinalis*, consists of all the pore-forming isoforms of CatSper (CatSper1 – 4) similar to mammals (24).

As CatSper is a complex protein with many subunits, it has not yet been successfully expressed functionally in cultured cells (18). There is only one study in which a voltage sensor domain of CatSper3 was expressed in the HEK293T cells and *Xenopus* oocytes (26). Therefore, the functional analysis of CatSper is performed using the spermatozoa from gene-deficient animals, with most studies being conducted in mice. In this study, we examined the role of CatSper to identify the key player involved in the influx of Ca^2+^ during sperm chemotaxis and also investigated the specific roles of CatSper in ascidians. Interestingly, the expression of CatSper in the ascidian sperm was not restricted to the sperm flagellum, but was also observed in the sperm head and other cells. Furthermore, we developed a *Catsper3*-deficient ascidian sperm using the CRISPR/Cas9 system and found that CatSper is also an indispensable Ca^2+^ channel in the ascidian sperm. Therefore, CatSper plays significant roles in both sperm chemotaxis and motility. CatSper may also play important roles in the development of sperms and spermatogenesis.

## Results

### Expression of CatSper in the ascidian

Initially, we examined the expression of CatSper in the ascidian *C. intestinalis* using RT-PCR. Surprisingly, in adults, all Catsper subtypes (*Catsper1, 2, 3, 4*) were expressed not only in the testis, but also in the heart (Figure 1A). Expression of the *Catsper* genes was also observed in the stomach and intestine; however, this may be due to the fact that testicular tissue exists in a form that is closely attached to the surface of the stomach and intestines, making it difficult to completely eliminate the testes during tissue collection. Furthermore, *Catsper3* and *Catsper4* were expressed in the siphon and gill. When the expression of Catsper was examined in the developmental juveniles, *Catsper3* and *Catsper4* were expressed in 2.5-week-old juveniles, although the expression was weak (Figure 1B). The 2.5-week-old juvenile is still immature, and the testes do not develop in this stage. Thus, the expression of *Catsper* in the ascidian does not seem to be restricted to the testis. On the other hand, by whole-mount *in situ* hybridization (WISH), *Catsper* expression was not detected in juveniles and was only observed in adult testes (Figure 2). These results indicate that the *Catsper* genes of the ascidian are mainly expressed in the testis, similar to those of mammalian species, but they are weakly expressed at places other than the testis.

**Fig. 1.**
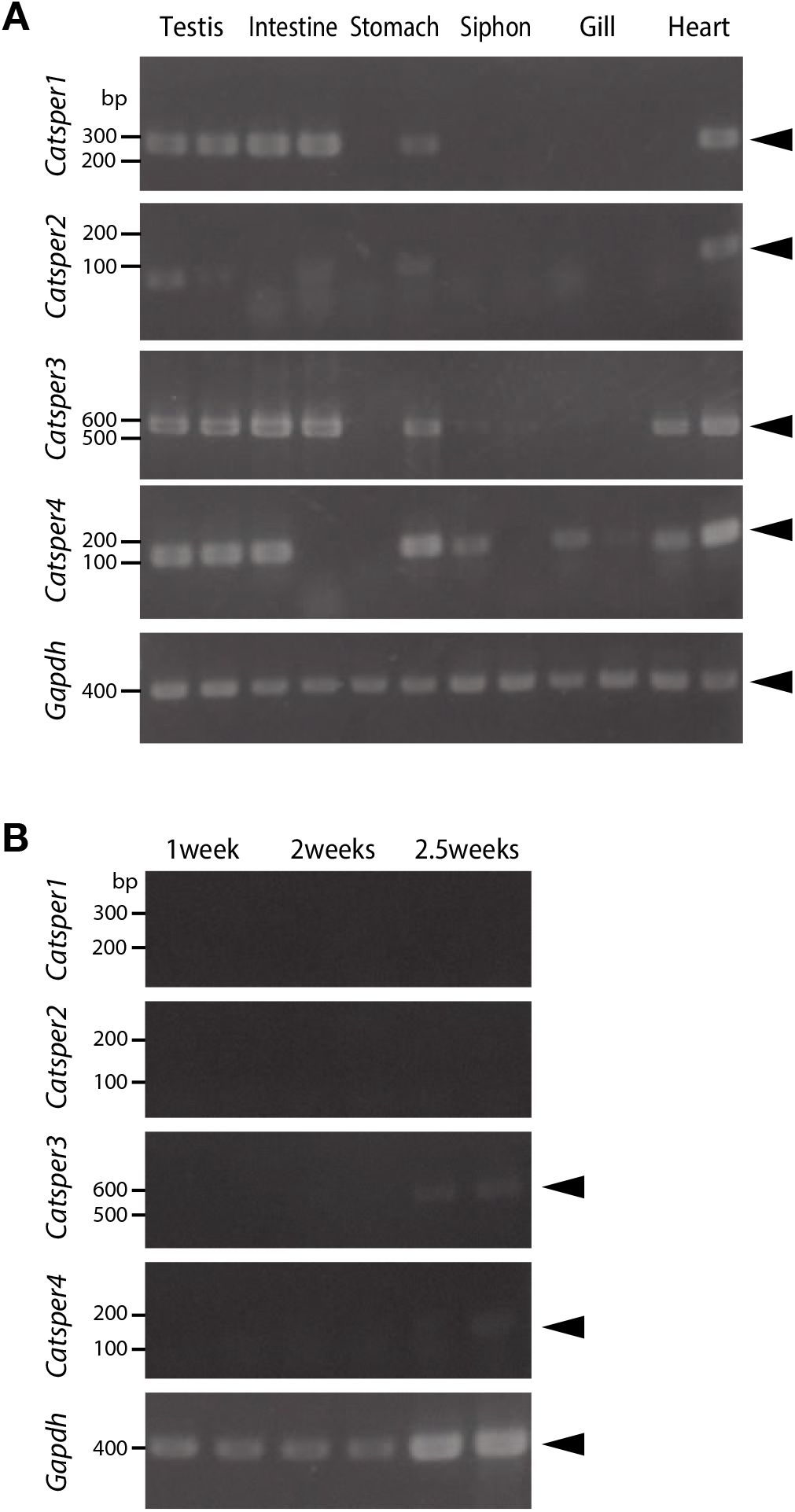
Expression levels of the *Catsper* genes in the ascidian, *Ciona intestinalis*, in the tissues of mature adults (A) and developing juveniles (B). Expression levels of the household gene, glyceraldehyde-3-phosphate dehydrogenase (*Gapdh*), was used as the control. Gene expression was examined by RT-PCR. Arrowheads indicate the bands pertaining to the products of RT-PCR.

**Fig. 2.**
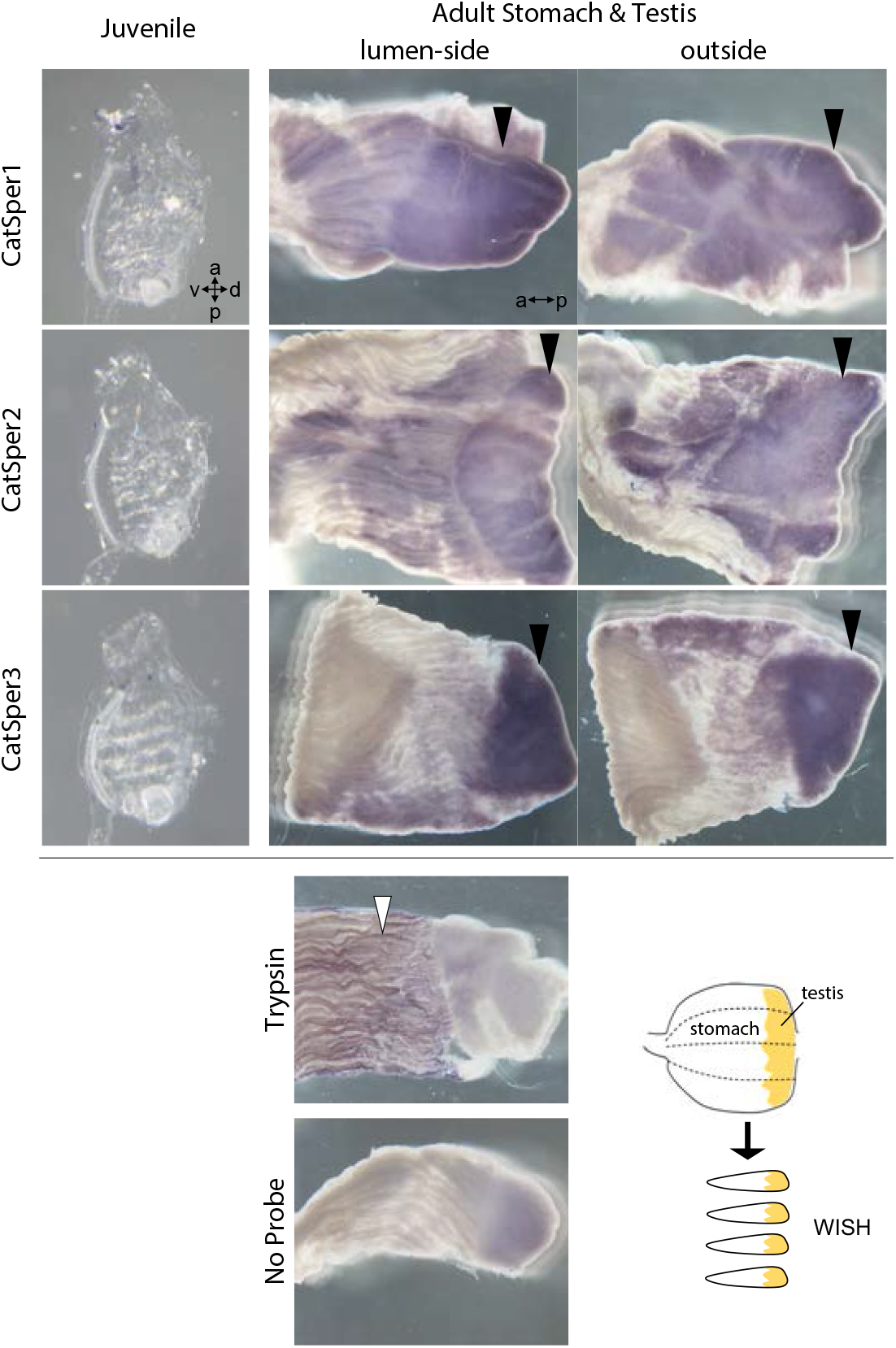
The whole mount in situ hybridization (WISH) method was used to analyze the expression levels of *Catsper1 Catsper2*, and *Catsper3* in the stomach and testes of the ascidian juveniles and adults. None of the *Catsper* subtypes were expressed in the juveniles. They were only expressed in the adult testes (black arrowhead). For positive control of WISH, the probe of *Trypsin-like* (<u>XP_002126930.1</u>) was used (white arrowhead).

### CatSper 3 was localized around the mitochondria of the ascidian sperm

Next, we examined the existence of CatSper proteins. As described above, the ascidian *Catsper* genes were mainly expressed in the testis, but were also weakly expressed in the heart, siphon, and gill. Thus, we examined whether CatSper proteins were present in these tissues. In mammalian species, CatSper proteins were observed only in the sperm flagella. On the other hand, in the ascidian, CatSper 3 protein was highly expressed in sperm (Figure 3). Interestingly, immunohistochemistry showed that CatSper3 seems to be expressed in the spermatogonia and head region of the sperm in the seminiferous tubules, and in the Leydig cell-like interstitial cells outside of the seminiferous tubules (Figure 4, upper and middle row). Precise observation of the sperm showed that CatSper 3 was localized not only in flagella but also around the mitochondria in the ascidian sperm head (Figure 5).

**Fig. 3.**
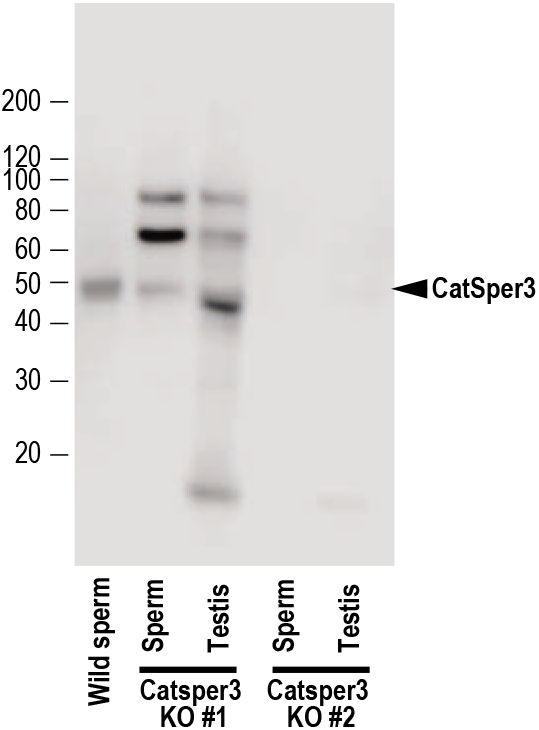
Representative results of the western blotting of the sperms and testes from two *Catsper3* knockout (KO) animals using the anti-CatSper3 antibody. The wildtype ascidian CatSper3 protein showed a 50 kDa band (arrowhead), while the predicted size from the deduced amino-acid sequence was 49809 Da.

**Fig. 4.**
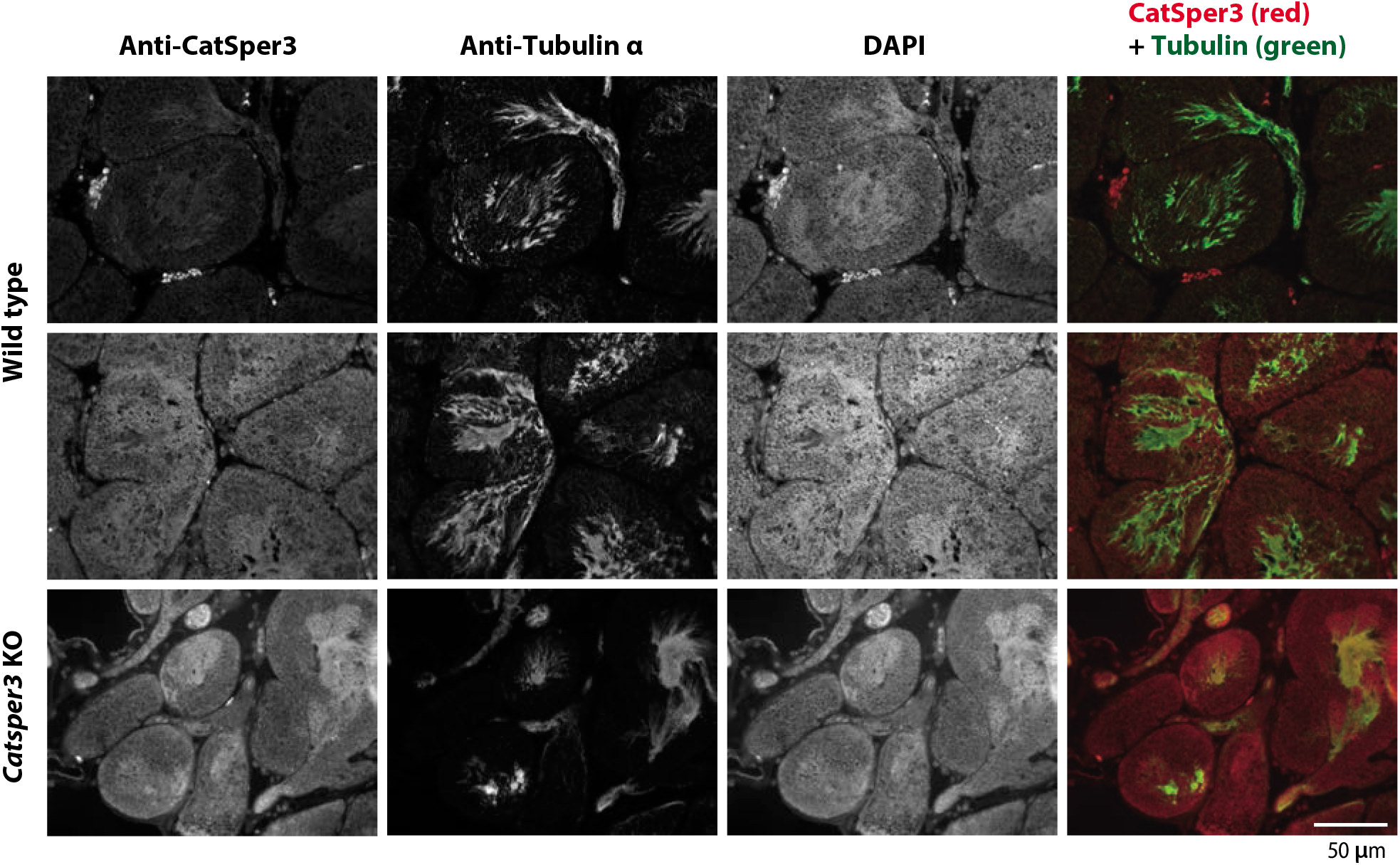
Immunohistchemistry with the anti-CatSper3 antibody in the ascidian testes. CatSper seems to be localized at the Leydig cells in the developing sperm cells. Scale bar = 50 μm.

**Fig. 5.**
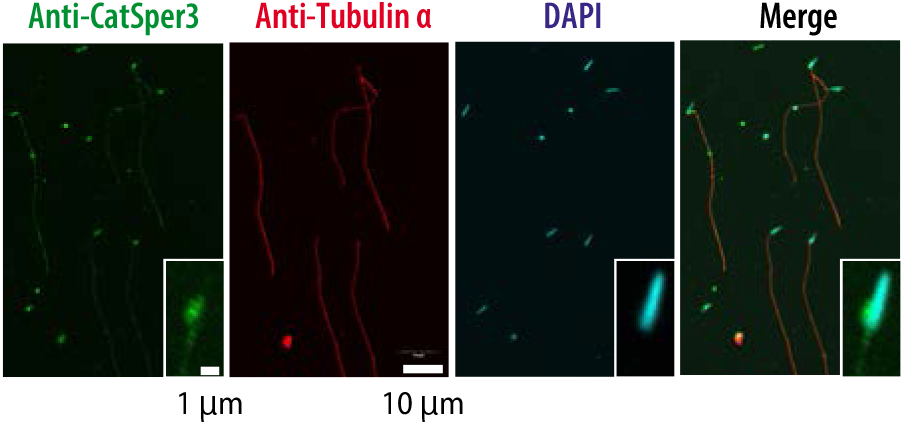
Indirect immunofluorescence assay with the anti-CatSper3 antibody in the ascidian sperm. (inset) Higher magnified image of the sperm head. The CatSper protein seems to be localized at the mitochondrial region of the head and also at the flagellum. Scale bar = 10 μm or 1 μm (insets).

### Development is delayed in the Catsper3 knockout animals

To examine the function of the CatSper channel in the ascidian sperm, we tried to produce a *Catsper-deficient* ascidian by using the CRISPR/Cas9 system. We selected exon 5 of *Catsper3* as the target gene of the CRISPR/Cas9 system (Figure 6A), because we could not find an effective target site for the CRISPR/Cas9 system on the other ascidian Catsper genes. The mutation frequency of the *Catsper3* gene by the CRISPR/Cas9 system was 94.7% (n = 38) when checked at the juvenile stage.

**Fig. 6.**
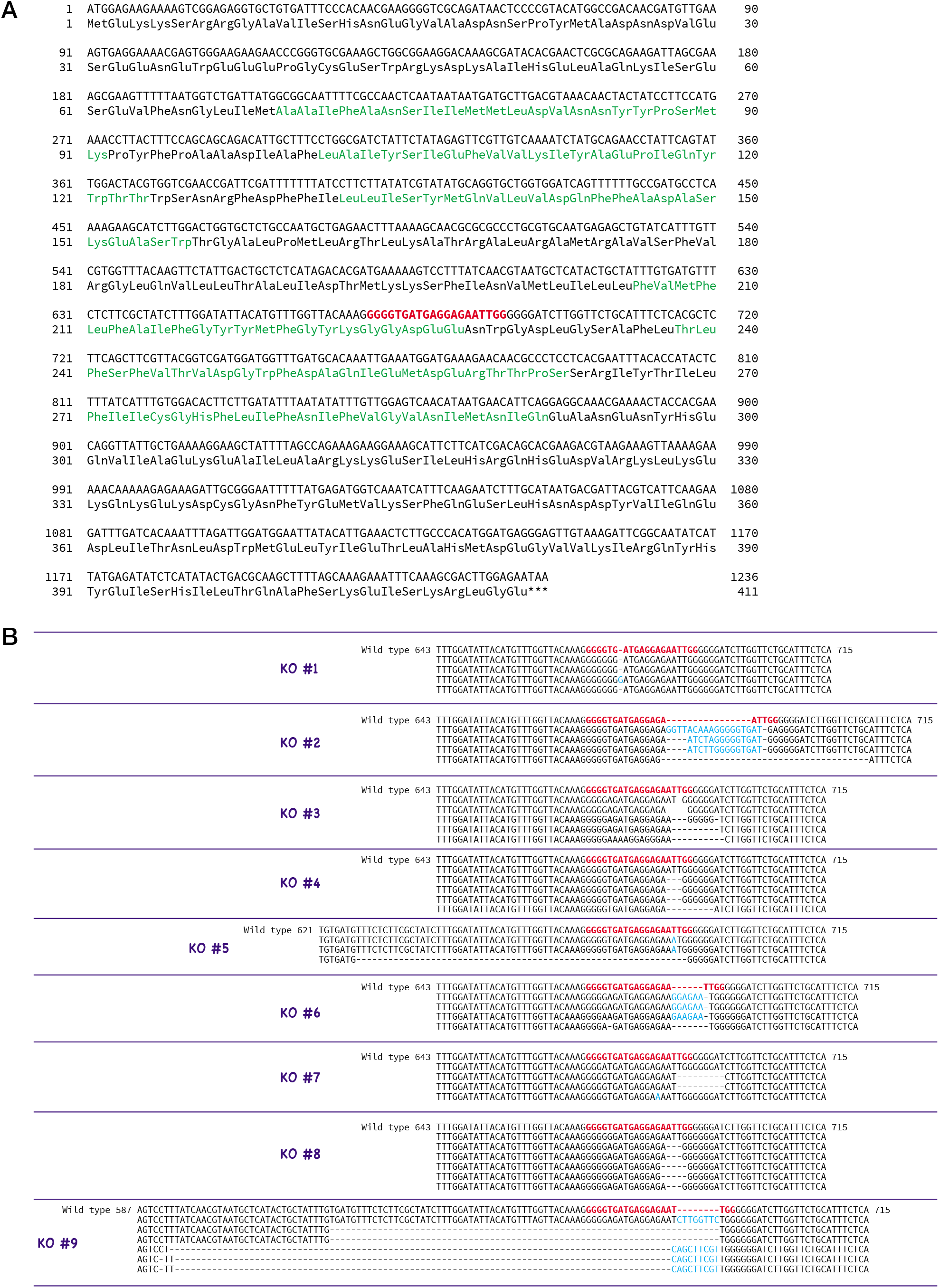
Generation of *Catsper3* KO ascidians. (A) DNA and amino acid sequences of *Catsper3*. Target sequence for the CRISPR/Cas9 system is shown in red bold letters. (B) Representative DNA sequences around the target sequence of the *Catser3* KO ascidians with the CRISPR/Cas9 system. The DNA sequence of un-muted wildtype *Catsper3* is shown in the upper row (wild type). Bold red letters show the target sequence for the CRISPR/Cas9 system. Deleted and inserted nucleotides are shown by a dash and blue letter, respectively.

Unfortunately, the F_0_ animals in which *the Catsper3* gene was edited by the CRISPR/Cas9 system (Catsper3 KO animals) tended to die when their tadpole larvae settled and metamorphosed into juveniles. Furthermore, the growth rate of the Catsper3 KO animals was much slower than that of the control; the body size of the two-month-old individuals was significantly smaller than that of the wild-type animals (Figure 7). Finally, we obtained nine KO animals that reached the reproductively mature stage (Figure 6B). Since the gametes from the KO animals could not fertilize, we used the KO animals for the following analyses.

**Fig. 7.**
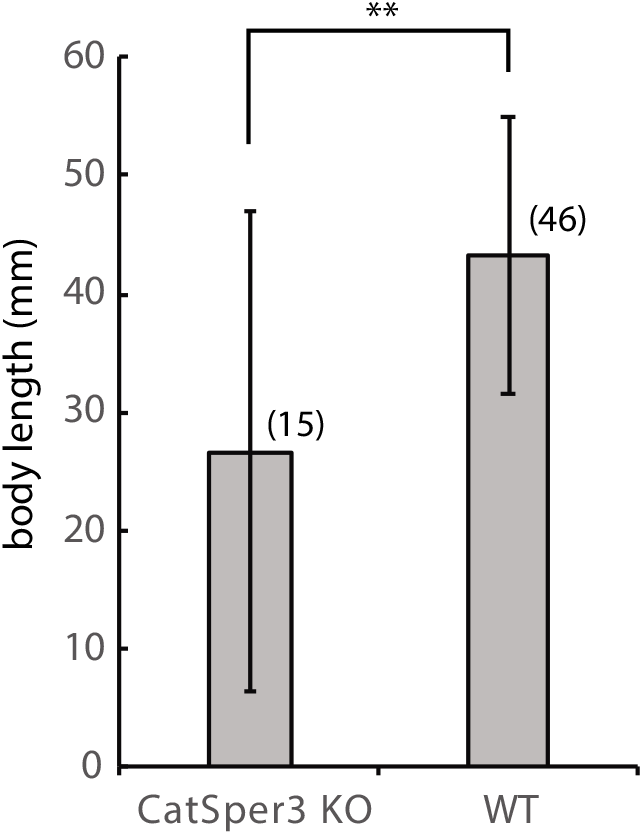
Body sizes of the *Catsper3* KO and wildtype (WT) animals. The examined KO and WT animals were 60–74 days-old and 47–75 days-old, respectively. Values are expressed as the mean ± standard deviation (S.D). The number in parenthesis represents the number of examined animals. There was a significant difference between the wildtype and KO animals as determined by the Student’s t-test. ** *P* < 0.01.

### Spermatogenesis of the Catsper3 knockout animals

In the sperm obtained from the CatSper3 KO animals, CatSper3 protein was still observed, and immunohistochemistry using the anti-CatSper3 antibody showed that CatSper3 was localized at the sperm head in the seminiferous tubule (Figure 4, lower row). However, the protein seemed to be modified: the antibody-reacted band in the western blotting was seen at 90 and 70 keDa, which was higher than the wild-type CatSper3 protein and 45 kDa, which was lower than the wild-type CatSper3 protein (Figure 3C). Furthermore, the signal of CatSper3 in Leydig-cell-like interstitial cells was stronger than that in spermatozoa (arrowhead).

The testes of CatSper3 KO animals were smaller than those of wild-type animals at the same age, which is consistent with the poor development of the individuals. Spermatogenesis seemed to occur normally in the KO animals (Figure 8, upper and middle row), which yielded mature spermatozoa whose shape appeared to be normal. However, spermatozoa from KO animals were fragile; headless or broken-flagella spermatozoa were often observed by microscopy (Figure 8, lower row).

**Fig. 8.**
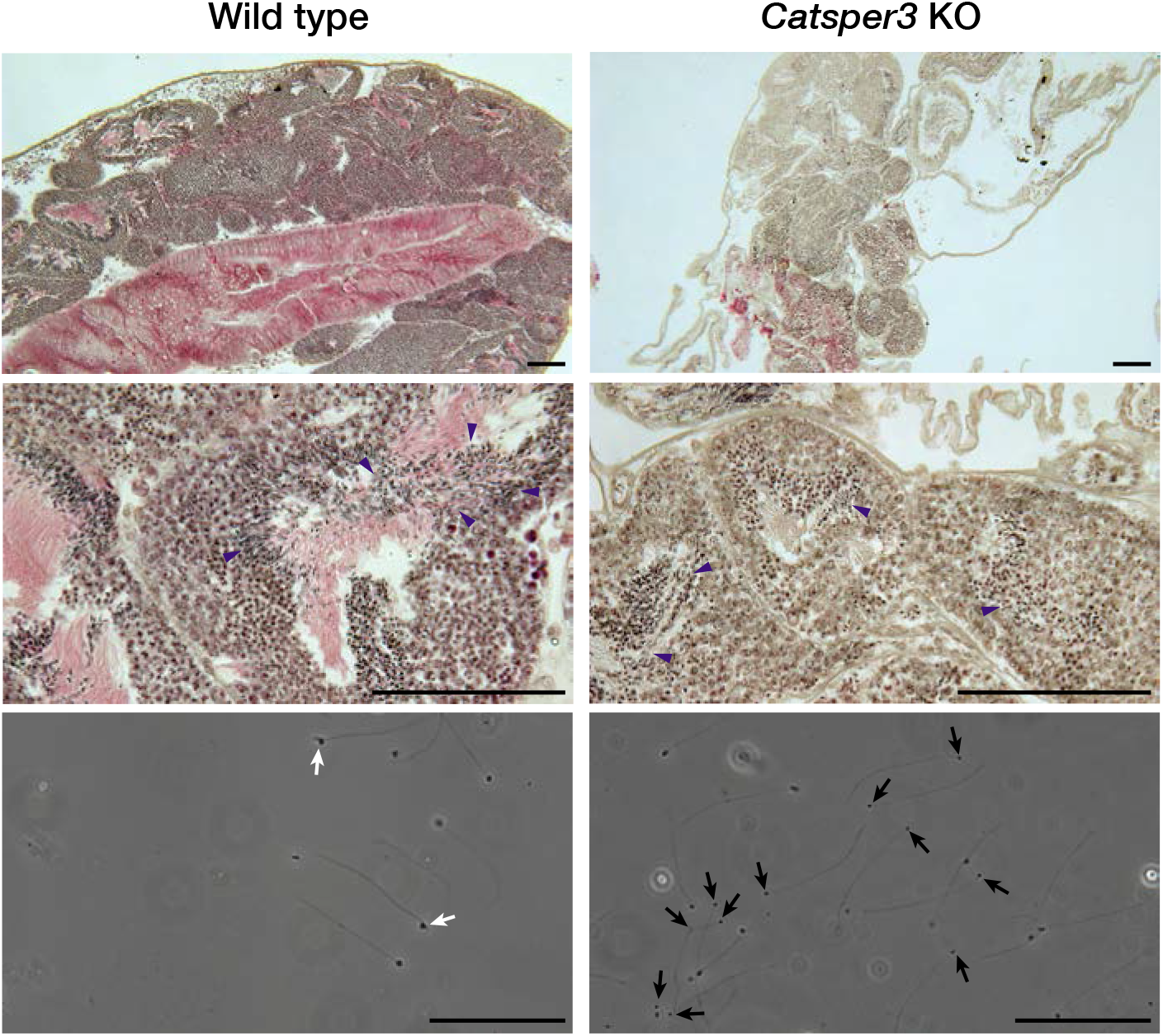
Histological assessment of the testes and sperm of the *Catsper3* KO ascidians. Lower (upper row) and higher (middle row) magnification of the testicular section taken from wildtype and *Catsper3* KO ascidians and stained with hematoxylin and eosin. Arrowheads indicate the sperms produced in the testis. (Lower row) Spermatozoa obtained from the spermiducts of the wildtype and *Catsper3* KO ascidians. Some spermatozoa of the KO animal lost their head or had broken flagellum (black arrows). Typical heads of the wildtype spermatozoa are indicated by white arrows. Scale bar = 50 μm.

### Sperm motility and chemotactic behavior of the Catsper3 knockdown animals

As described above, the spermatozoa from the KO animals looked fragile and headless or broken-flagella spermatozoa were often observed (Figure 8, lower row). The sperm appeared normal in shape, and almost all sperm had no motility even in the presence of theophylline, which is an activating agent of ascidian sperm motility (Supplemental movies S1 and S2). Few spermatozoa showed forward motility, and those seemed to have a large lateral amplitude of the head (Supplemental movie S2, Figure 9), similar to sperm movement in the presence of PMCA inhibitor (11). In addition, none of the spermatozoa showed a chemotactic response (Figure 9).

**Fig. 9.**
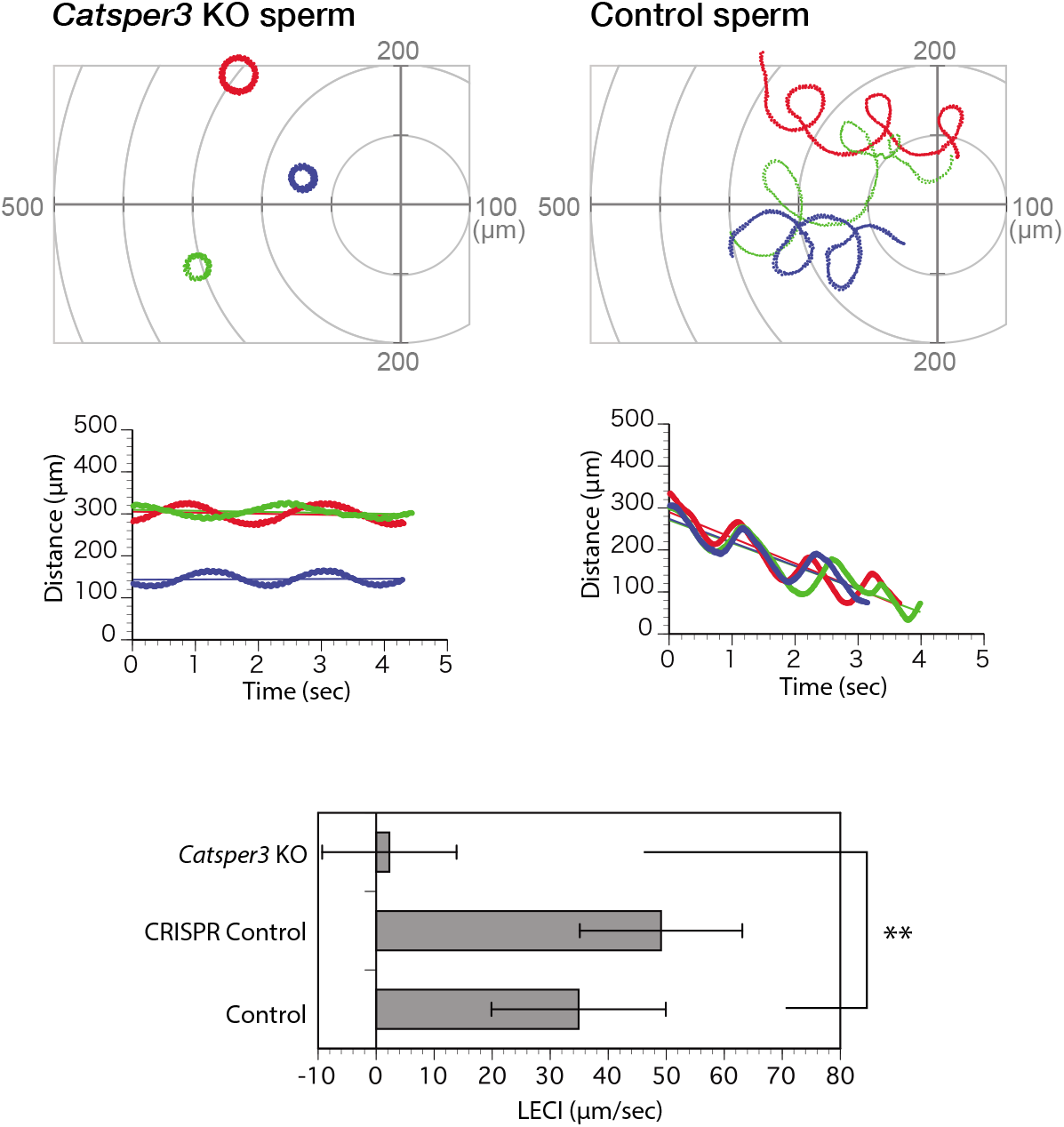
Chemotactic behavior of the Catsper3-knockout sperm. (Upper row) Typical three trajectories of the sperms of *Catsper3* KO and wildtype animals suspended in seawater around the tip of a capillary containing 5 μM sperm-activating and -attracting factor (SAAF). The origin of the coordinates indicates the capillary tip. (Middle row) Changes in the distance between the capillary tip and the head of the swimming sperm. The lines represent the linear equation of time vs. distance. The chemotaxis index (LECI) is calculated as negative values of the coefficients in the equation. (Lower row) Chemotaxis index (LECI) of the sperm. CRISPR control is the data about the sperm from an animal who was injected with Cas9 and gRNA but did not show any genetic mutation (n = 1). Values are expressed as the mean ± S.D. Statistical significance is set at \*\**P* < 0.01 (Student’s t-test).

## Discussion

In the 20 years since the discovery of CatSper as a sperm-specific Ca^2+^ channel (12), many studies have shown that CatSper is the only important Ca^2+^ channel that functions in sperm in mammals (27). CatSper is localized in linear quadrilateral nanodomains along the flagellum in mammalian sperm and mediates rheotactic behavior in the female reproductive tract (18,28). On the other hand, many animals, including birds and amphibians, lack the CatSper gene (24) and it is not known whether CatSper has the same functions in sperm of animals other than mammals, although only in the sea urchin CatSper seems to mediate chemotactic behavior in pharmacological studies (29). The ascidian *C. intestinalis* belongs to chordates and has four isoforms of CatSper (CatSper1 – 4), similar to mammals. Interestingly, CatSper seems to be not restricted to sperm, and our study showed that expression of CatSper in the ascidian was not restricted to the testis, but it was also observed in non-mature juvenile and adult hearts, even though strong expression of CatSper genes was observed in the testis (Figure 1). The distribution of CatSper in the ascidian sperm also differs from that in mammalian sperm: in the ascidian sperm, CatSper3 is located at the flagellum and mitochondrial region in the head (Figure 5). Furthermore, most of the Catsper3-KO ascidians died within a month because of stunted growth, and the body size of the mature Catsper3-KO ascidians was significantly small. These data suggest that CatSper plays a role in the development and/or nutrient uptake. Even though the expression area of CatSper3 in organs other than the testis is still not examined, cilia seemed to be immunoreactive to the anti-CatSper3 antibody. Thus, CatSper might be expressed in the cilia and flagella not only in the sperm but also in the whole body, and may play a role in the movement of cilia/flagella.

The shape of the sperm obtained from the Catsper3-KO ascidians was almost normal, but that from some animals looked abnormal, with a small head, and they were fragile: the head was easily disconnected from the flagellum (Figure 8). Furthermore, the spermatozoa were almost quiescent and never activated by any stimulations. Some sperm had motility, but the sperm did not show chemotactic behavior. In mice, there are no morphological differences between wild-type and CatSper-KO sperm in any subunit (12,15–19,30). On the other hand, spermatogenesis in the Catsper3-KO male was not different from that in the wild type (Figure 8). Whether the morphological abnormalities in spermatozoa are due to abnormal spermatogenesis or low nutrition caused by eating disorders will require further detailed analysis in the future.

The CatSper-KO sperm did not show any response to SAAF, and no chemotactic behavior was observed. This result is consistent with the pharmacological findings on sperm chemotaxis in the sea urchin *Arbacia punctulata* (25). In mammals, sperm lacking CatSper are motile but cannot show hyperactivation, resulting in infertility (12,15). Therefore, it is plausible that CatSper is involved in the regulation of flagellar motility in ascidians, sea urchins, and mammals. In mammals, CatSper is assumed to be a direct or indirect target of various chemicals and is involved in chemotaxis (31). In humans, progesterone, which is a putative sperm attractant of mammals, has been shown to activate CatSper by binding to α/β hydrolase domain-containing protein 2 (32). Furthermore, subunits of CatSper, EFCAB9, and Catsperζ regulate the opening of CatSper channels in response to changes in intracellular Ca^2+^ and pH (19). On the other hand, in the ascidian, the sperm attractant SAAF directly binds to PMCA and seems to induce Ca^2+^ efflux from the sperm (11). Furthermore, in the ascidian, the CatSper-deficient sperm not only lacked chemotactic behavior, but also had significant effects on motility itself, with many sperm failing to initiate the movement. In mammals, sperm lacking any pore-forming CatSper subunit have similar initial velocities as wild-type (15), and CatSperζ-null and EFCAB9-null spermatozoa are also motile, although they show abnormal flagellar waveforms (18,19). Thus, the system regulating sperm flagellar beating, including the CatSper channel, seems to differ from mammals, even though the system is mainly regulated by [Ca^2+^]_i_: CatSper is also an indispensable Ca^2+^ channel in the ascidian sperm, but it plays a role not only in sperm flagellar beatings but also in fundamental sperm motility.

In conclusion, the results of this study indicate that CatSper plays an important role in the ascidian sperm. Unlike in mammals, CatSper seems to be involved not only in the regulation of the sperm flagellar beatings, but also in the sperm motility in ascidians. Moreover, the expression of CatSper in the ascidian was not restricted to the sperm flagellum and was also observed in non-mature juveniles, adult hearts, Leydig cells, and sperm heads. CatSper may also play a role in the development of sperms and spermatogenesis. Although CatSper seems to play more significant roles in ascidians than mammals, it may only be due to impaired spermatogenesis; however, further studies are needed to confirm this. We also aim to further explore the different roles of CatSper in future studies.

## Experimental procedures

### Materials

The ascidian *C. intestinalis* (type A; also called *C. robusta*) was obtained from the National BioResource Project for *Ciona* (http://marinebio.nbrp.jp/). The ascidian sperm was obtained as described previously (33). Artificial seawater contained 462 mM NaCl, 9 mM KCl, 10 mM CaCl2, 48 mM MgCl2, and 10 mM HEPES-NaOH (pH 8.2). SAAF and its derivatives were synthesized as described previously (9,34). Juveniles were developed by artificial insemination using mature eggs and the sperm obtained from dissected gonoducts. The specimens of whole-mount juveniles and dissected adult specimens were prepared for in situ hybridization as described by Nakayama and Ogasawara (35).

Reagents not specifically mentioned in the text were purchased from FUJIFILM Wako Pure Chemicals (Osaka, Japan).

### RT-PCR

Predicted *Catsper* genes were identified from the genome database of *C. intestinalis* (http://ghost.zool.kyoto-u.ac.jp/cgi-bin/gb2/gbrowse/kh/). Total RNA was extracted from each specimen using the RNeasy Mini Kit (Qiagen, Hilden, Germany), and the concentration of RNA was determined by absorbance at 260 nm in relation to the absorbance at 280 nm. One microgram of total RNA was reverse transcribed to cDNA using a Transcriptor First Strand cDNA Synthesis Kit (Roche Diagnostics, Tokyo Japan) using an anchored-oligo(dT)18 primer and random hexamer primer. The 50 μL reaction mixture contained 32 μL nuclease-free water, 1.0 μL cDNA, 1.5 μL (10 μmol/L) of each primer, 5 μL (2 mM) dNTPs, 3 μL (25 mM) MgSO4, 5 μL (10x) KOD plus ver.2 buffer, and 1 μL KOD plus (TOYOBO, Osaka, Japan). The primers used were as follows: *Catsper1* (KH.C10.377): 5’-TCACACAGCATGGAC GGTAA-’3 and 5’-GCATCAGGATTGCTCCTAAAGA-’3; *Catsper2* (KH.S391.7): 5’-CGCCCAAGTATCATCCTGAA-’3 and 5’-GACATGTACTTAATAGTGGCAACACC-’3; *Catsper3* (KH. C2.323): 5’-CGAAAGCGAAGTTTTTAATGGTCT-’3 and 5’-GCACTTCCTTATAAGCCATGTGTAA-’3; *Catsper4* (KH.C2.993): 5’-AGGAAGAACAGCGGTCGAAG-’3 and 5’-CTGAGGCTTCAGAGCATCCA-’3; and *Gapdh:* 5’-ACCCAGAAGACAGTGGATGG-’3 and 5’-CAGGACACCAGCTTCACAAA-’3. The amplification cycle was as follows: denaturation at 94 °C for 2 min, followed by 30 cycles of denaturation at 98 °C for 10 s, annealing at 55 °C for 30 s, and at 68 °C for 1 min. PCR products were analyzed on 2% (wt/vol) agarose gels stained with 0.5 μg/mL ethidium bromide and visualized under UV light. Images of the gels were captured using a gel imaging device (Printgraph, AE-6932; ATTO, Tokyo, Japan), and the acquired images were processed using Adobe Photoshop and Adobe Illustrator (Adobe Systems, San Jose, CA, USA).

### Whole-mount in situ hybridization

Digoxigenin-labeled antisense RNA probes for the *Ciona* Catsper genes were synthesized from T7-RNA polymerase promoter-attached amplified cDNA, and the probes were purified by centrifugal ultrafiltration as described previously (36). WISH was performed on the *Ciona* specimens using “InSitu Chip,” as described by Ogasawara et al. (37). Gene expression signals were visualized using nitroblue tetrazolium/5-bromo-4-chloro-3-indolylphosphate solutions using a standard method (Roche). Whole-mount and sectioned specimens of *Ciona* were observed using an SZX12 stereo microscope (Olympus, Tokyo, Japan).

### Gene knockdown by CRISPR/Cas9 system

The CRISPR/Cas9 method (38,39) was used for genome editing in *C. intestinalis*, as described previously (40). We designed the target sequences of gRNA that are specific for *Catsper3* (5’-CGAAAGCGAAGTTTTTAATGGTCT-3’) at the fifth exon, using ZiFiT (http://zifit.partners.org/ZiFiT) (41,42). The target DNAs were inserted into the BsaI site of pDR274 (Addgene; 39) as vectors for *in vitro* transcription of gRNAs. The *Cas9* gene, which was a gift from George Church (hCas9) (Addgene plasmid #41815; http://n2t.net/addgene:41815; RRID:Addgene_41815) (43) was subcloned into pCS2+ as a vector for *in vitro* transcription of mRNAs. gRNAs and mRNAs of Cas9 were synthesized using MEGAshortscript T7 and MEGAshortscript SP6 Transcription Kits (Thermo Fisher Scientific, Waltham, MA USA), respectively.

We injected approximately 30 to 100 pL of the RNA solution containing 400 ng/μL gRNA, 800 ng/μL Cas9 mRNA, and 10% phenol red to unfertilized eggs, and reared embryos and animals in aquariums.

We extracted DNA from the juveniles or muscles of adults and carried out PCR to determine the efficiency of the genome editing using the heteroduplex mobility assay (44). The primers used for PCR were: 5’-TTGACTGCTCTCATAGACACGATG-3’ and 5’-TTACCGTAACGAAGCTGAAGAGC-3’. After purification of the PCR bands by electrophoresis, the PCR products were subcloned into pGEM-T Easy (Promega, Madison, WI USA) for sequencing analyses using a Genetic Analyzer (ABI 3130; Thermo Fisher Scientific).

### Western blotting

The samples were directly solubilized in NuPAGE LDS sample buffer (Novex, Carlsbad, CA, USA). The solubilized samples were treated with benzonase nuclease (Novagen, Billerica, MA, USA) and the proteins were separated by NuPAGE SDS-PAGE Gel System using 3–8% Tris-Acetate Gels (Novex) and transferred to PVDF membranes. The anti-CatSper3 antibody (ab197924; Abcam, Tokyo, Japan), horseradish peroxidase (HRP)-conjugated anti-rabbit IgG antibody (GE Healthcare Japan, Tokyo, Japan), and ECL Prime (GE Healthcare Japan) were used for the detection of CatSper. Luminescence was imaged using a gel imager (EzCapture II; ATTO). The acquired images were processed using Photoshop and Illustrator (Adobe, San Jose, CA USA).

### Indirect immunofluorescence

The sperm suspension was placed onto a coverslip, fixed with 4% paraformaldehyde for 10 min, and washed twice with ASW for 5 min. The fixed sperm were permeabilized with 0.1% NP-40 for 15 min, blocked with 1% BSA for 1 h, and incubated with 0.6 μg/mL anti-α tubulin monoclonal antibody (#236-10501; Thermo Fisher Scientific) or 4.3 μg/mL anti-CatSper3 antibody with 1% BSA in PBS overnight. The spermatozoa were washed three times with PBS and incubated with x1/2000 Alexa 488-conjugated anti-mouse IgG (H+L) (Thermo Fisher Scientific) or x1/2000 DyLight550-conjugated anti-rabbit IgG (Abcam) for 30 min. After washing twice with PBS, the spermatozoa were observed and acquired using a confocal laser-scanning microscope (FV3000; Olympus).

### Immunohistochemistry

The testes were dissected and fixed in Bouin’s fixative consisting of 71.4% saturated picric acid, 23.8% formalin, and 4.7% acetic acid, for 2 h at room temperature. The tissues were embedded in paraffin and cut into 4 μm-thick sections. Some sections were stained with hematoxylin and eosin for morphological observation and made into permanent specimens. For immunohistochemical analysis, the antigens were retrieved by heating in a citrate buffer (pH 6.0). The sections were incubated with 10% goat serum (Nichirei, Tokyo, Japan) as a blocking solution, and incubated with 0.6 μg/mL anti-α tubulin monochronal antibody (#236-10501; Thermo Fisher Scientific) or 4.3 μg/mL anti-CatSper3 antibody with 10% goat serum (Nichirei) overnight. Then, the sections were washed and incubated with x1/2000 Alexa 488-conjugated anti-mouse IgG(H+L) (Thermo Fisher Scientific) or x1/500, Alexa Fluor 546-conjugated anti-rabbit IgG (H+L) (Thermo Fisher Scientific) for 1 h. After washing three times with PBS, the sections were mounted with ProLong™ Gold Antifade Mountant with DAPI (Thermo Fisher Scientific). Tissue sections were observed using a fluorescent microscope (IX-71; Olympus). Images were recorded on a PC connected to a digital CCD camera (Retiga Exi; QImaging, Surrey, Canada) using an imaging application (TI workbench) (45). The acquired data were processed using Adobe Photoshop and Adobe Illustrator (Adobe).

### Analysis of sperm motility and chemotaxis

Sperm chemotaxis was examined as described previously (5). Briefly, the semen was diluted 10^4^–10^5^ times in ASW with 1 mM theophylline (Sigma-Aldrich Japan, Tokyo, Japan) to induce the activation of motility (7). The activated-sperm suspension was placed into the observation chamber, and sperm movement around the micropipette tip containing the SAAF was recorded. The position of the sperm head was determined using Bohboh software (BohbohSoft, Tokyo, Japan) (46). The parameters of chemotactic activity, including the trajectory, distance between the capillary tip and sperm, and LECI were calculated as described previously (8).

### Statistical analysis

All experiments were repeated at least three times with different specimens. Data are expressed as the mean ± SD. Statistical significance in Fig. 7 and 9 was calculated using the Student’s *t*-test. P < 0.01 was considered to be statistically significant.

## Data availability

All data are contained in the manuscript.

## Supporting information

This article contains supporting information.

## Acknowledgments

We would like to thank Prof. T. Miura for allowing us to use CLSM. We also thank Mr. H. Kohtsuka, Ms. N. Ito, Ms. M Kawabata, and Ms. M. Kohtsuka (MMBS, University of Tokyo) for their technical assistance. We also thank Mr. S. Nakayama for his technical assistance with *in situ* hybridization. We would also like to thank the National Bio-Resource Project of the Japan Agency for Medical Research and Development (AMED) for supplying the materials used in this study.

## Funding

This work was supported by a grant to MY from the Japan Society for the Promotion of Science (JSPS) KAKENHI (20K06715).

## Conflict of interest

The authors declare that they have no conflicts of interest with the contents of this article.

## Abbreviations

CatSper: cation channel of sperm
KO: knockout
LECI: liner equation-based chemotaxis index
PMCA: plasma-menbrane type Ca^2+^/ATPase
SAAF: sperm-activation and attaction factor
WISH: whole-mount *in situ* hybridization

## Supporting information

**Supplementary movie S1.** Behavior of the wild-type sperm around the capillary containing 1 μM sperm-activating and -attracting factor (SAAF).

**Supplementary movie S2.** Behavior of the *Catsper3* knockout (KO) sperm around the capillary containing 1 μM SAAF.

## References

1. Yoshida, M., and Yoshida, K. (2018) Modulation of Sperm Motility and Function Prior to Fertilization. in Reproductive and Developmental Strategies. Diversity and Commonality in Animals. (Kobayashi, K., Kitano, T., Iwao, Y., and Kondo, M. eds.), Springer, Tokyo. pp 437–462

2. Yoshida, M., and Yoshida, K. (2011) Sperm chemotaxis and regulation of flagellar movement by Ca2+. Mol. Hum. Reprod. 17, 457–465

3. Brokaw, C. J., Josslin, R., and Bobrow, L. (1974) Calcium ion regulation of flagellar beat symmetry in reactivated sea urchin spermatozoa. Biochem. Biophys. Res. Commun. 58, 795–800

4. Brokaw, C. J. (1979) Calcium-induced asymmetrical beating of triton-demembranated sea urchin sperm flagella. J. Cell Biol. 82, 401–411

5. Shiba, K., Baba, S. A., Inoue, T., and Yoshida, M. (2008) Ca2+bursts occur around a local minimal concentration of attractant and trigger sperm chemotactic response. Proc Natl Acad Sci USA 105, 19312–19317

6. Yoshida, M., Ishikawa, M., Izumi, H., De Santis, R., and Morisawa, M. (2003) Store-operated calcium channel regulates the chemotactic behavior of ascidian sperm. Proceedings of National Academy Sciences USA 100, 149–154

7. Yoshida, M., Inaba, K., Ishida, K., and Morisawa, M. (1994) Calcium and cyclic-AMP mediate sperm activation, but Ca2+alone contributes sperm chemotaxis in the ascidian, Ciona savignyi. Dev. Growth Diff 36, 589–595

8. Yoshida, M., Murata, M., Inaba, K., and Morisawa, M. (2002) A chemoattractant for ascidian spermatozoa is a sulfated steroid. Proceedings of National Academy Sciences USA 99, 14831–14836

9. Oishi, T., Tsuchikawa, H., Murata, M., Yoshida, M., and Morisawa, M. (2004) Synthesis and identification of an endogenous sperm activating and attracting factor isolated from eggs of the ascidian *Ciona intestinalis*; an example of nanomolar-level structure elucidation of novel natural compound. Tetrahedron 60, 6971–6980

10. Miyashiro, D., Shiba, K., Miyashita, T., Baba, S. A., Yoshida, M., and Kamimura, S. (2015) Chemotactic response with a constant delay-time mechanism in Ciona spermatozoa revealed by a high time resolution analysis of flagellar motility. Biol. Open 4, 109–118

11. Yoshida, K., Shiba, K., Sakamoto, A., Ikenaga, J., Matsunaga, S., Inaba, K., and Yoshida, M. (2018) Ca2+efflux via plasma membrane Ca2+-ATPase mediates chemotaxis in ascidian sperm. Sci Rep 8, 16622

12. Ren, D., Navarro, B., Perez, G., Jackson, A. C., Hsu, S., Shi, Q., Tilly, J. L., and Clapham, D. E. (2001) A sperm ion channel required for sperm motility and male fertility. Nature 413, 603–609

13. Bystroff, C. (2018) Intramembranal disulfide cross-linking elucidates the super-quaternary structure of mammalian CatSpers. Reprod Biol 18, 76–82

14. Lishko, P. V., Kirichok, Y., Ren, D., Navarro, B., Chung, J. J., and Clapham, D. E. (2012) The control of male fertility by spermatozoan ion channels. Annu. Rev. Physiol. 74, 453–475

15. Qi, H., Moran, M. M., Navarro, B., Chong, J. A., Krapivinsky, G., Krapivinsky, L., Kirichok, Y., Ramsey, I. S., Quill, T. A., and Clapham, D. E. (2007) All four CatSper ion channel proteins are required for male fertility and sperm cell hyperactivated motility. Proc. Natl. Acad. Sci. USA 104, 1219–1223

16. Liu, J., Xia, J., Cho, K. H., Clapham, D. E., and Ren, D. (2007) CatSperβ, a novel transmembrane protein in the CatSper channel complex. J. Biol. Chem. 282, 18945–18952

17. Chung, J. J., Navarro, B., Krapivinsky, G., Krapivinsky, L., and Clapham, D. E. (2011) A novel gene required for male fertility and functional CATSPER channel formation in spermatozoa. Nat. Commun. 2, 153

18. Chung, J. J., Miki, K., Kim, D., Shim, S. H., Shi, H. F., Hwang, J. Y., Cai, X., Iseri, Y., Zhuang, X., and Clapham, D. E. (2017) CatSperzeta regulates the structural continuity of sperm Ca2+ signaling domains and is required for normal fertility. Elife 6

19. Hwang, J. Y., Mannowetz, N., Zhang, Y., Everley, R. A., Gygi, S. P., Bewersdorf, J., Lishko, P. V., and Chung, J. J. (2019) Dual Sensing of Physiologic pH and Calcium by EFCAB9 Regulates Sperm Motility. Cell 177, 1480–1494 e1419

20. Carlson, A. E., Westenbroek, R. E., Quill, T., Ren, D., Clapham, D. E., Hille, B., Garbers, D. L., and Babcock, D. F. (2003) CatSper1 required for evoked Ca2+entry and control of flagellar function in sperm. Proc. Natl. Acad. Sci. USA 100, 14864–14868

21. Quill, T. A., Ren, D., Clapham, D. E., and Garbers, D. L. (2001) A voltage-gated ion channel expressed specifically in spermatozoa. Proceedings of National Academy Sciences USA 98, 12527–12531

22. Zeng, X. H., Navarro, B., Xia, X. M., Clapham, D. E., and Lingle, C. J. (2013) Simultaneous knockout of Slo3 and CatSper1 abolishes all alkalization- and voltage-activated current in mouse spermatozoa. J. Gen. Physiol. 142, 305–313

23. Schiffer, C., Rieger, S., Brenker, C., Young, S., Hamzeh, H., Wachten, D., Tuttelmann, F., Ropke, A., Kaupp, U. B., Wang, T., Wagner, A., Krallmann, C., Kliesch, S., Fallnich, C., and Strunker, T. (2020) Rotational motion and rheotaxis of human sperm do not require functional CatSper channels and transmembrane Ca(2+) signaling. EMBO J 39, e102363

24. Cai, X., and Clapham, D. E. (2008) Evolutionary genomics reveals lineage-specific gene loss and rapid evolution of a sperm-specific ion channel complex: CatSpers and CatSperbeta. PLoS One 3, e3569

25. Seifert, R., Flick, M., Bonigk, W., Alvarez, L., Trotschel, C., Poetsch, A., Muller, A., Goodwin, N., Pelzer, P., Kashikar, N. D., Kremmer, E., Jikeli, J., Timmermann, B., Kuhl, H., Fridman, D., Windler, F., Kaupp, U. B., and Strunker, T. (2015) The CatSper channel controls chemosensation in sea urchin sperm. The EMBO Journal 34, 379–392

26. Arima, H., Tsutsui, H., Sakamoto, A., Yoshida, M., and Okamura, Y. (2018) Induction of divalent cation permeability by heterologous expression of a voltage sensor domain. Biochim Biophys Acta 1860, 981–990

27. Lishko, P. V., and Mannowetz, N. (2018) CatSper: A Unique Calcium Channel of the Sperm Flagellum. Curr Opin Physiol 2, 109–113

28. Chung, J. J., Shim, S. H., Everley, R. A., Gygi, S. P., Zhuang, X., and Clapham, D. E. (2014) Structurally distinct Ca2+signaling domains of sperm flagella orchestrate tyrosine phosphorylation and motility. Cell 157, 808–822

29. Rennhack, A., Schiffer, C., Brenker, C., Fridman, D., Nitao, E. T., Cheng, Y. M., Tamburrino, L., Balbach, M., Stolting, G., Berger, T. K., Kierzek, M., Alvarez, L., Wachten, D., Zeng, X. H., Baldi, E., Publicover, S. J., Benjamin Kaupp, U., and Strunker, T. (2018) A novel cross-species inhibitor to study the function of CatSper Ca(2+) channels in sperm. Br J Pharmacol 175, 3144–3161

30. Quill, T. A., Sugden, S. A., Rossi, K. L., Doolittle, L. K., Hammer, R. E., and Garbers, D. L. (2003) Hyperactivated sperm motility driven by CatSper2 is required for fertilization. Proc. Natl. Acad. Sci. USA 100, 14869–148674

31. Brenker, C., Goodwin, N., Weyand, I., Kashikar, N. D., Naruse, M., Krahling, M., Muller, A., Kaupp, U. B., and Strunker, T. (2012) The CatSper channel: a polymodal chemosensor in human sperm. The EMBO Journal 31, 1654–1665

32. Miller, M. R., Mannowetz, N., Iavarone, A. T., Safavi, R., Gracheva, E. O., Smith, J. F., Hill, R. Z., Bautista, D. M., Kirichok, Y., and Lishko, P. V. (2016) Unconventional endocannabinoid signaling governs sperm activation via the sex hormone progesterone. Science 352, 555–559

33. Yoshida, M., Inaba, K., and Morisawa, M. (1993) Sperm chemotaxis during the process of fertilization in the ascidians *Ciona savignyi* and *Ciona intestinalis*. Dev. Biol. 157, 497–506

34. Oishi, T., Tuchikawa, H., Murata, M., Yoshida, M., and Morisawa, M. (2003) Synthesis of endogenous sperm-activating and attracting factor isolated from ascidian *Ciona intestinalis*. Tetrahedron lett. 44, 6387–6389

35. Nakayama, S., and Ogasawara, M. (2017) Compartmentalized expression patterns of pancreatic- and gastric-related genes in the alimentary canal of the ascidian Ciona intestinalis: evolutionary insights into the functional regionality of the gastrointestinal tract in Olfactores. Cell and tissue research 370, 113–128

36. Ogasawara, M., Minokawa, T., Sasakura, Y., Nishida, H., and Makabe, K. W. (2001) A large-scale whole-mount in situ hybridization system: Rapid one-tube preparation of DIG-labeled RNA probes and high throughput hybridization using 96-well silent screen plates. Zool. Sci. 18, 187–193

37. Ogasawara, M., Satoh, N., Shimada, Y., Wang, Z., Tanaka, T., and Noji, S. (2006) Rapid and stable buffer exchange system using InSitu Chip suitable for multicolor and large-scale whole-mount analyses. Dev Genes Evol 216, 100–104

38. Cong, L., Ran, F. A., Cox, D., Lin, S., Barretto, R., Habib, N., Hsu, P. D., Wu, X., Jiang, W., Marraffini, L. A., and Zhang, F. (2013) Multiplex genome engineering using CRISPR/Cas systems. Science 339, 819–823

39. Hwang, W. Y., Fu, Y., Reyon, D., Maeder, M. L., Tsai, S. Q., Sander, J. D., Peterson, R. T., Yeh, J. R., and Joung, J. K. (2013) Efficient genome editing in zebrafish using a CRISPR-Cas system. Nat. Biotechnol. 31, 227–229

40. Sasaki, H., Yoshida, K., Hozumi, A., and Sasakura, Y. (2014) CRISPR/Cas9-mediated gene knockout in the ascidian *Ciona intestinalis*. Dev. Growth Differ. 56, 499–510

41. Sander, J. D., Maeder, M. L., Reyon, D., Voytas, D. F., Joung, J. K., and Dobbs, D. (2010) ZiFiT (Zinc Finger Targeter): an updated zinc finger engineering tool. Nucleic Acids Res. 38, W462–468

42. Sander, J. D., Zaback, P., Joung, J. K., Voytas, D. F., and Dobbs, D. (2007) Zinc Finger Targeter (ZiFiT): an engineered zinc finger/target site design tool. Nucleic Acids Res. 35, W599–605

43. Mali, P., Yang, L., Esvelt, K. M., Aach, J., Guell, M., DiCarlo, J. E., Norville, J. E., and Church, G. M. (2013) RNA-guided human genome engineering via Cas9. Science 339, 823–826

44. Ota, S., Hisano, Y., Muraki, M., Hoshijima, K., Dahlem, T. J., Grunwald, D. J., Okada, Y., and Kawahara, A. (2013) Efficient identification of TALEN-mediated genome modifications using heteroduplex mobility assays. Genes Cells 18, 450–458

45. Inoue, T. (2018) TI Workbench, an integrated software package for electrophysiology and imaging. Microscopy (Oxf) 67, 129–143

46. Shiba, K., Mogami, Y., and Baba, S. A. (2002) Ciliary Movement of Sea-urchin Embryos. Natural Science Report, Ochanomizu Univ. 53, 49–54

